# PolyJet 3D Printing of Tissue Mimicking Materials: An Investigation of Characteristic Properties of 3D Printed Synthetic Tissue

**DOI:** 10.1101/2020.12.23.424075

**Authors:** Vania Lee, Leah Severseike, Chris Bakken, Emily Bermel, Varun Bhatia

## Abstract

Current anatomical 3D printing has been primarily used for education, training, and surgical planning purposes. This is largely due to the models being printed in materials which excel at replicating macro-level organic geometries; however, these materials have the drawback of unrealistic mechanical behavior and system properties compared to biological tissue. The new Digital Anatomy (DA) family of materials from Stratasys utilizes composite printed materials to more closely mimic mechanical behavior of biological tissue, potentially allowing more realistic models for design evaluation. Various experimental DA Solid Organ (SO) configurations were quantitatively evaluated under axial loading for comparison with porcine liver in terms of stiffness. Additionally, Structural Heart - Myocardium (Myo) configurations were quantitatively evaluated under different lubricant conditions for comparison with porcine epicardium and aorta in terms of lubricity. Finally, experimental DA Subcutaneous Tissue configurations were qualitatively evaluated by experts with significant pre-clinical implant experience for cutting, tunneling, and puncture procedures.

In general, the experimental SO configurations showed promising compliance results when compared to porcine liver. The stiffness of DA configurations was either within the same range or on the lower bound of porcine tissue stiffness values. The lubricity of DA configurations with surface treatments was comparable with porcine epicardium and aorta. In terms of qualitative cutting, DA did not perform well for any of the configurations; however, tunneling and puncture were rated favorably for some of the experimental configurations. Despite some limitations, DA feels closer to real tissue than other commercially available 3D printed materials. Furthermore, the lower sample-to-sample variability of DA allows for repeatability not provided by biological tissue. The promising results and repeatability indicate that DA materials can be used to configure structures with similar characteristic mechanical properties to porcine liver, epicardium, and subcutaneous tissue, adding new value as not only an educational, training, and surgical tool, but also as a research tool.

## Introduction

Anatomical 3D printing has been primarily used for education, training, and surgical planning purposes [1–5], with limited success in quantitative evaluations [6–8]. This is primarily due to the limitations of 3D printing materials not being able to mimic the material and mechanical properties of tissue. Anatomical models which mimic tissue both in macro structure as well as mechanical characteristics have the potential to bridge bench level ex vivo testing and in vivo animal or cadaver testing. Furthermore, models which can match the lubricity of biological tissue would provide realistic use conditions and haptics for procedural evaluations.

Use of phantoms or 3D printed anatomical models offer benefits compared to in-vivo and ex-vivo animal and cadaveric testing including reduced cost, longevity, and no need to meet ethical regulatory requirements. Additionally, the performance of a device can be confounded by unwanted (or unknown) variations in animal/cadaver tissue properties, resulting in longer product development times and extra cost. Agar, gelatin gel, and silicone are some of the tissue mimicking materials commonly used to develop anatomical models [9–11]; however, models created from these materials require conventional fabrication processes such as casting and molding, which are not optimal for creating patient-specific anatomy models in terms of process time and tooling costs. Use of 3D printing technologies address some of the limitations of conventional manufacturing processes, allowing for quicker and lower cost development of models for both idealized as well as patient-specific anatomies.

Currently, the material selection available for 3D printed models is not formulated with the intention of matching mechanical properties or lubricity of biological tissue. A new selection of Digital Anatomy (DA) materials developed by Stratasys (Eden Prairie, MN) has been designed to have varying degrees of stiffness based on the thickness of the outer Agilus shell (30A) and an infill mixture of TissueMatrix (Shore 00-30) and Agilus in specific infill patterns. These material-structure combinations include tissue mimicking material configurations such as myocardium, soft organ, and subcutaneous tissue.

The work presented in this paper is the latter part of a two-phase process of evaluating characteristics and mechanical properties of DA 3D printed materials through a combination of quantitative and qualitative assessments. While the previous work focused on the mechanical properties of DA Myocardium (Myo) [12], the work presented here is focused on (i) the mechanical properties of DA Soft Organ (SO) – an experimental configuration aimed at simulating liver tissue - compared to porcine liver, (ii) lubricity of DA Myo (highly contractile, very stiff, and extremely stiff) compared to porcine epicardium and aorta, and (iii) procedure specific tactile response of DA Subcutaneous Tissue compared to procedures conducted in porcine models.

Porcine tissue was chosen as the comparative animal model due to availability and common use in preclinical testing. The tests, and the corresponding results presented in this paper, were developed to mimic approximate boundary conditions of tools during in-vivo procedures. These included axial compression for stiffness testing (liver), friction/lubricity testing (epicardium and aorta), and qualitative cutting, tunneling and puncture testing (subcutaneous tissue) for comparison between DA materials and porcine tissue.

## Methods

### Soft Organ (Liver) Stiffness

#### Tissue Collection

Porcine liver was obtained from a pre-clinical research facility supplier in a recently harvested unfrozen state, shipped on ice. Once received, the gallbladder was removed and the liver was either wrapped in 1 x Phosphate-Buffered Saline (PBS) soaked paper towels and frozen in a −80°C freezer or immediately prepared for testing.

For sample preparation, livers were thawed just prior to use and only in testable quantities. Livers were thawed by placing in an insulated cooler overnight, allowing for about 12 to 16 hours of thaw time [9, 13]. Once thawed, livers were dissected by using a 3D printed cutter of dimensions 20 cm x 20 cm, creating samples with corresponding base dimensions as shown in Figure 1. The base dimensions were chosen to minimize test artifact from the force transferring member, a 3 mm diameter rounded rod, while allowing multiple sample specimens to be harvested from the porcine liver. Samples were taken from each lobe of the liver (Left Lateral, Left Medial, Right Medial, and Right Lateral) as shown in Figure 2. Due to the smaller size of the medial lobes, sample frequencies were lower for medial lobes compared to lateral lobes. All samples were tested within two hours of dissection.

**Figure 1.**
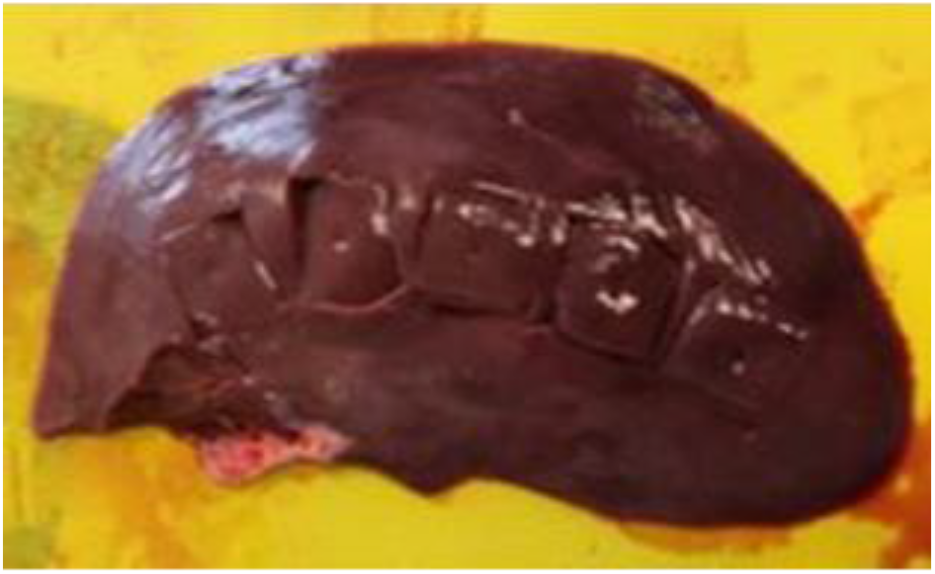
Example of dissected samples prepared from the left lateral lobe of a porcine liver

**Figure 2.**
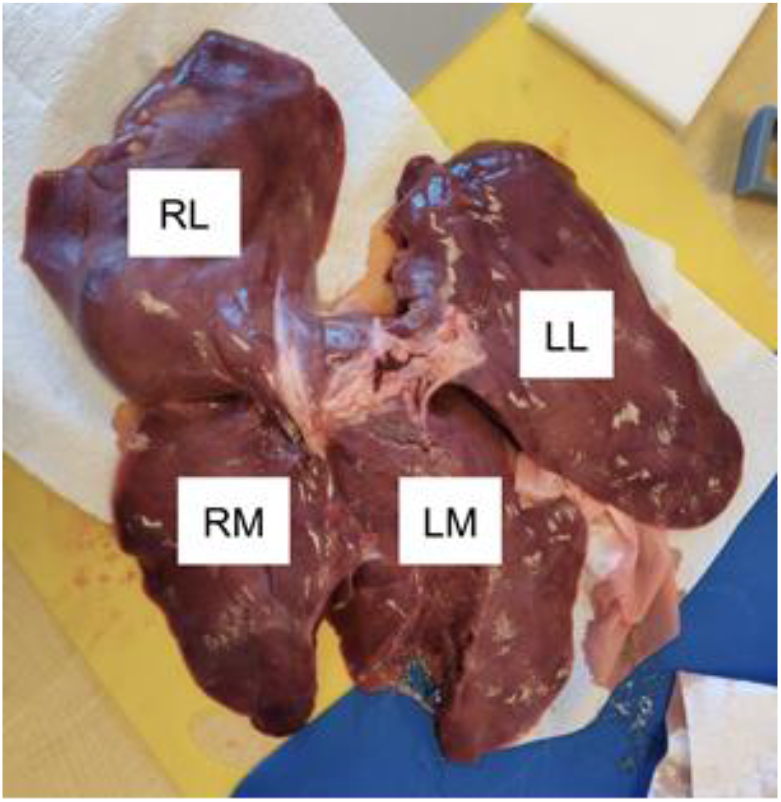
Tissue samples were taken from all four lobes of porcine liver: Right Lateral (RL), Right Medial (RM), Left Medial (LM), and Left Lateral (LL).

#### Stiffness Testing

Stiffness testing (axial compression) was performed on the 20 cm x 20 cm base tissue samples by placing them within a 3D printed base fixture providing wall and base support to the tissue. The boundary conditions were chosen to reflect use conditions for probing the liver during a procedure. The base fixture was mounted to the base of a single column Instron (equipped with 50 N load cell) and a rounded rod 3 mm in diameter was mounted to the top with a three-jaw chuck. The 3 mm diameter rod was selected to simulate a tool used during an implant procedure. The test method consisted of probing the constrained liver sample in axial compression with the rounded rod at a rate of 0.5 in/min, and the test setup is shown below in Figure 3 (left). The output was a force vs. displacement curve which was analyzed from 1 - 3 N to reflect realistic and safe use conditions. Larger forces would likely be outside the range expected in typical clinical use cases.

**Figure 3.**
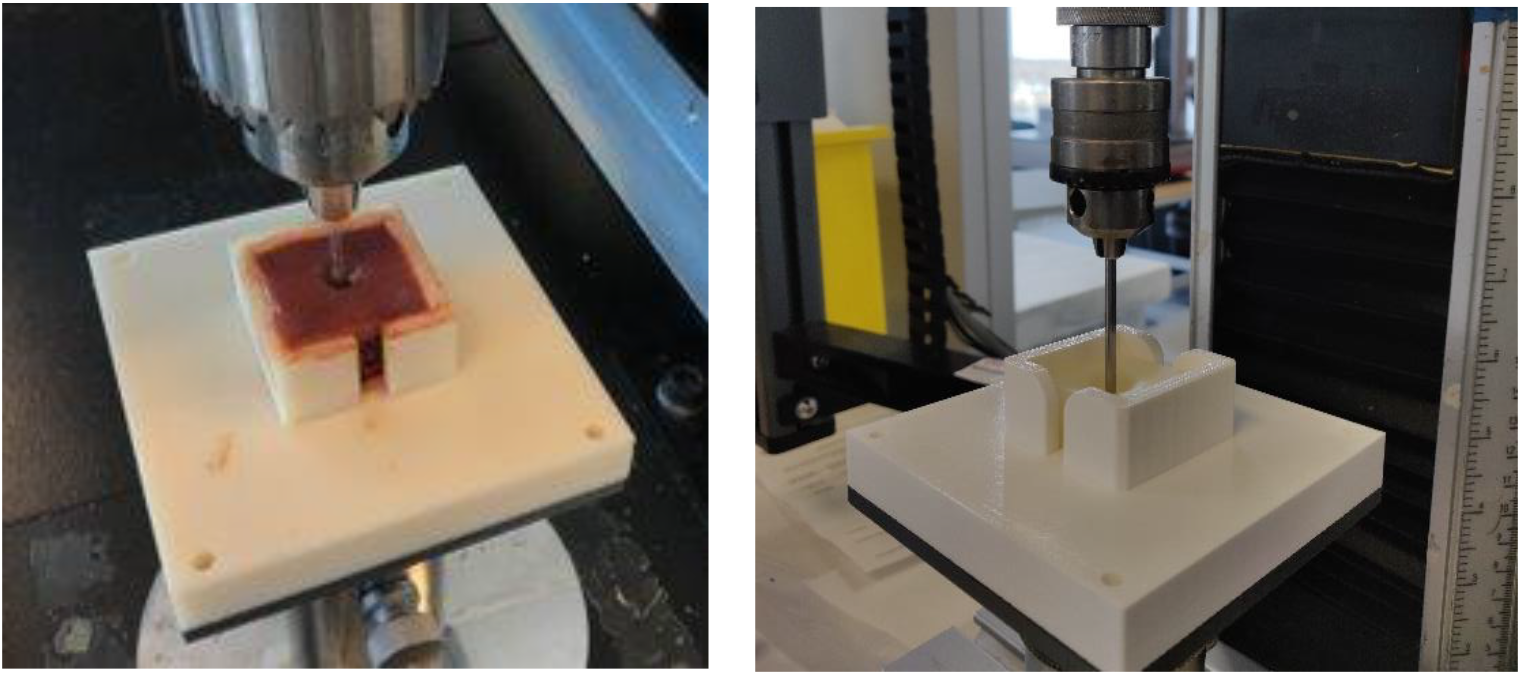
(Left): The liver tissue sample is constrained by the bottom 3D printed fixture. (Right): The DA liver sample is constrained in the bottom fixture. (Both): The 3mm diameter rounded rod is applying an axial compressive force, generating a force vs. displacement curve as the test output.

From the thickness values of the porcine liver samples, two representative thickness values were chosen as dimensions for the DA liver cube samples: 15 mm and 25 mm. Samples were tested in the same manner as the porcine liver samples, bounded by the 3D printed bottom fixture with an applied compressive load through a 3 mm diameter rounded rod. The test setup is shown in Figure 3 (right). Two iterative sets of liver configurations (Liver 1 - Liver 7) were tested, with replicate testing of Iteration 1: Liver 2 based on an initial filter test. Descriptions of the DA liver sample configurations are provided in Table 1.

**Table 1.** Tested configurations for DA SO Liver samples

| Sample Configurations | Agilus Shell [mm] |
| --- | --- |
| Iteration 1: Liver 2 | 0.4 |
| Iteration 2: Liver 1 – Liver 7 | 0.4 |
| Iteration 1: Liver 2 | 1.0 |
| Iteration 2: Liver 1 – Liver 7 | 1.0 |

### Lubricity (Epicardium and Aorta)

#### Tissue Collection

Porcine heart with aorta was obtained from a pre-clinical research facility supplier in a recently harvested unfrozen state, shipped on ice. Once received, the hearts were wrapped in 1xPBS soaked paper towels and frozen in a −80°C freezer.

For sample preparation, hearts were thawed just prior to use and only in testable quantities. Hearts were thawed by placing in an insulated cooler overnight, allowing for about 12 to 16 hours of thaw time. Once thawed, the ventricular chambers of the hearts were dissected into slabs of consistent height, creating level samples. Aortas were dissected open and laid flat. Dissected samples from the heart and aorta are shown in Figure 4.

**Figure 4.**
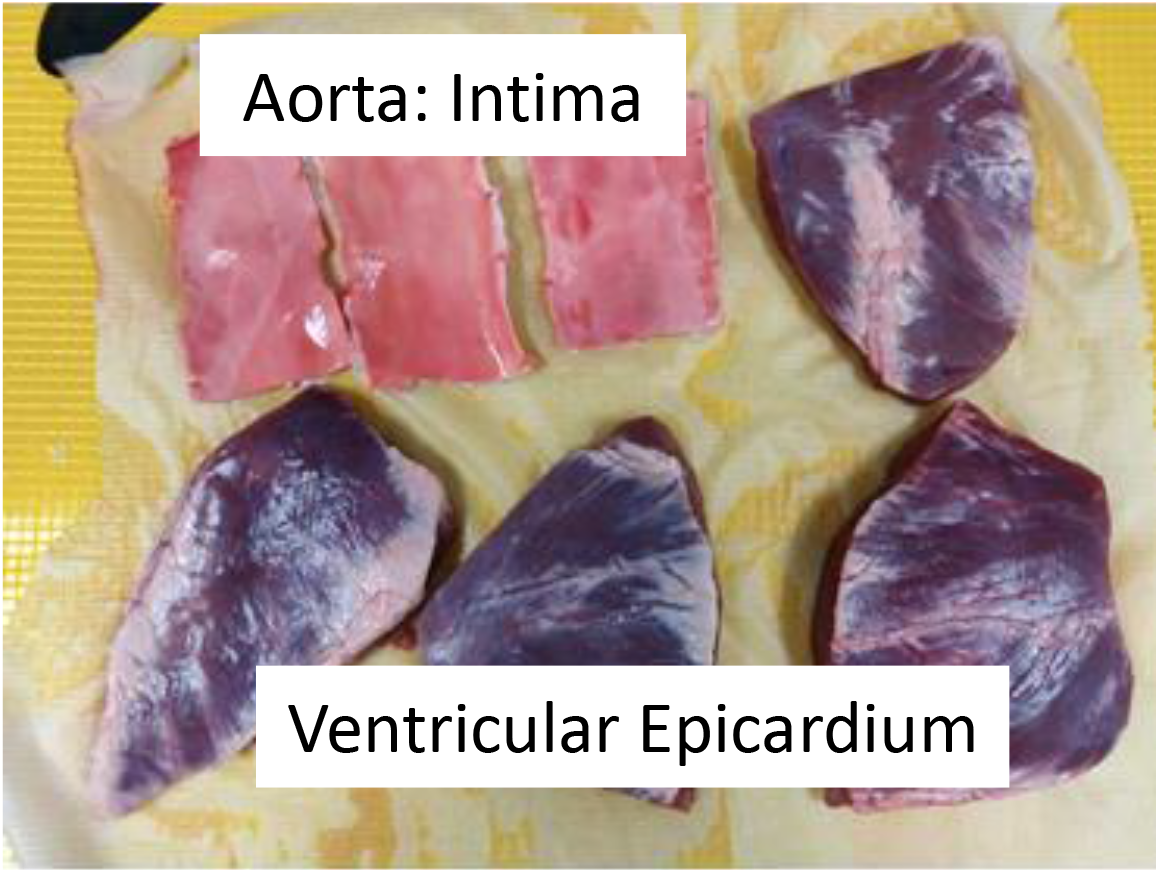
Dissected samples of aorta and ventricular chambers from a porcine heart

#### Lubricity Testing

For porcine tissue samples corresponding to DA Myo (highly contractile, very stiff, and extremely stiff), ventricular epicardium was chosen due to the lack of trabeculae. Furthermore, the intima of the aorta was chosen due to its surface interaction with delivery system tools. Porcine tissue samples were mounted epicardium or intima facing up to a 3D printed fixture lined with a silicone mat using T-pins. The assembly was then mounted to the base of a Rtec (San Jose, CA) tribometer. A 0.25 in diameter steel ball was used as the upper member of the wear couple, dragging along the surface of the mounted tissue with an applied axial force of 0.75 N at a velocity of 0.5 mm/s for a distance of 30 mm. These conditions were based upon preliminary testing to balance minimizing noise and plowing of the test sample while representing reasonable use conditions. The tissue was prevented from drying out by applying 1 x PBS between multiple runs. The test setup for the tissue samples are shown in Figure 5. Furthermore, the output was a coefficient of friction vs. time curve which was analyzed for a stabilized window of time.

**Figure 5.**
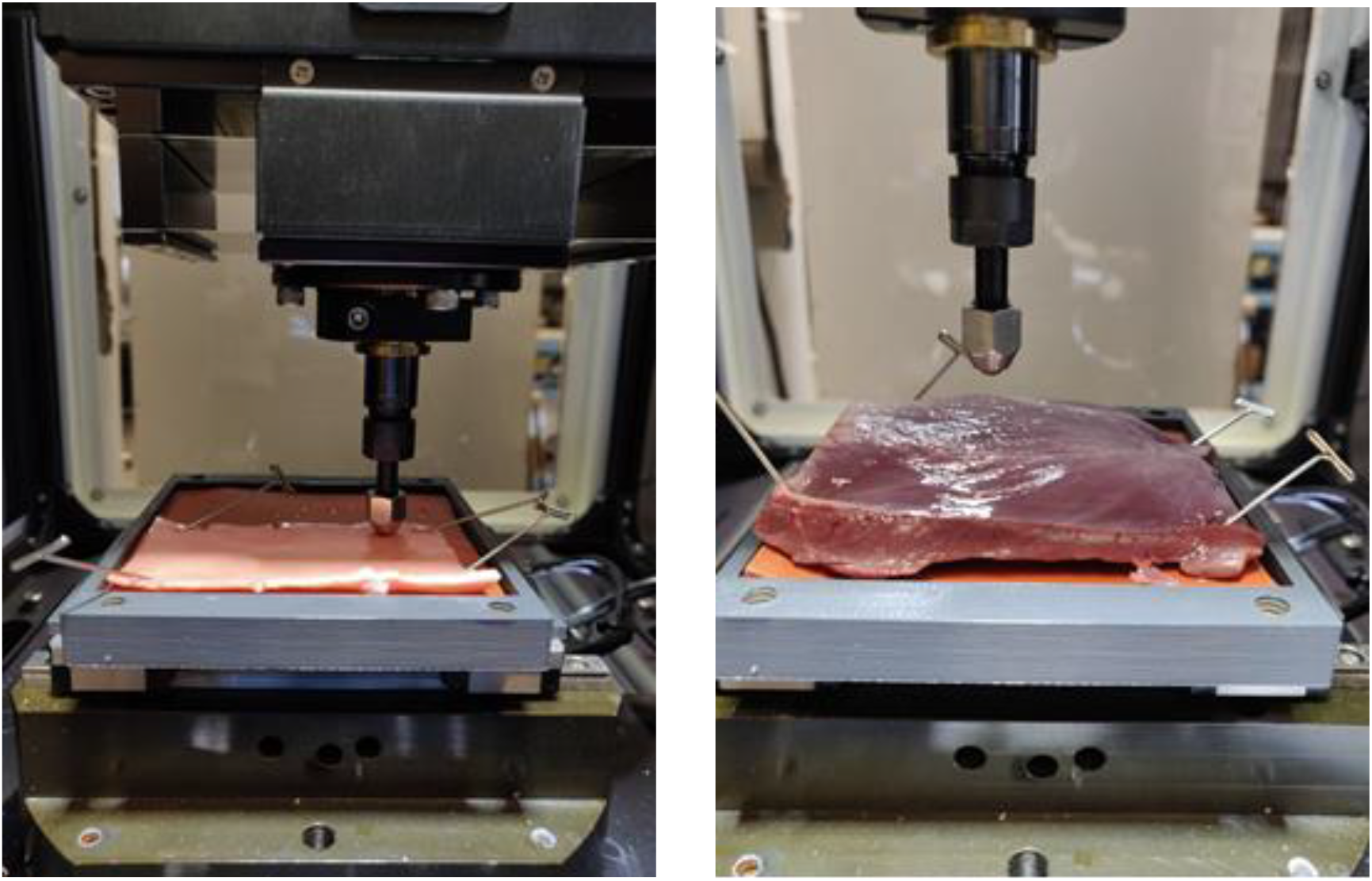
(Left): Porcine ventricle sample, epicardium side up, mounted to the silicone lined fixture with T-pins. (Right): Porcine aorta sample, intima side up, mounted to the silicone lined fixture with T-pins. (Both): 0.25 in diameter steel ball upper member as test probe.

Using the lower representative thickness values of porcine myocardial tissue, a thickness of 6 mm was chosen for the DA Myo samples. A rigid test fixture consisting of an inner insert of DA Myo (matte) was attached to the base of the tribometer and tested with a 0.25 in diameter steel ball probe with the following lubricant conditions: none (dry), DI water, mineral oil, and dish soap. Lubricants were chosen based upon cost effectiveness, availability, and ease of use. Figure 6 shows a representative mounted sample.

**Figure 6.**
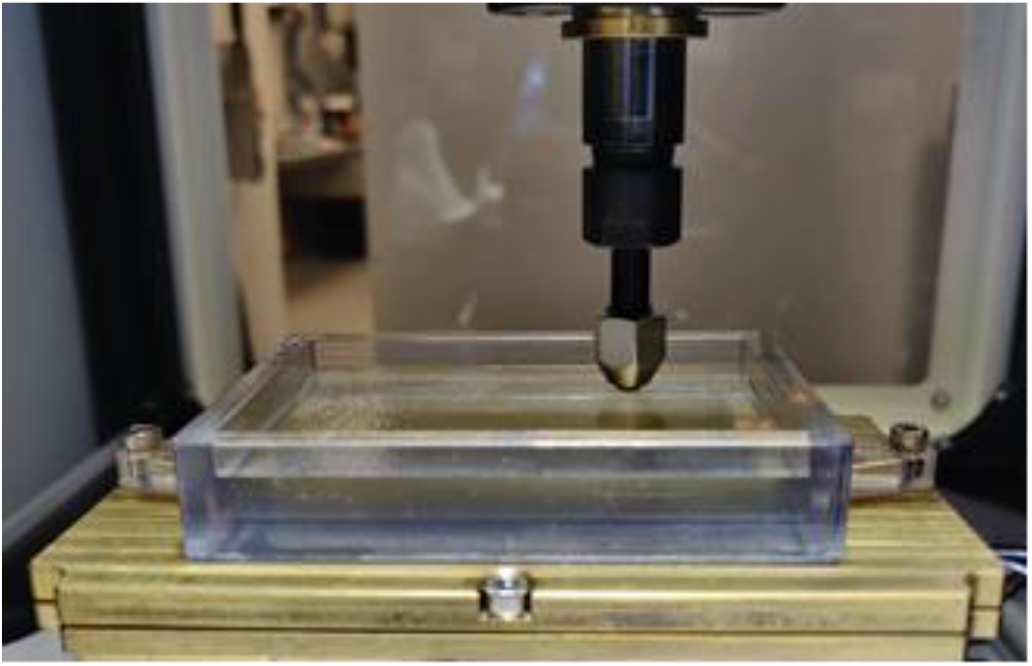
Rigid 3D printed fixture with DA Myo in the center with steel ball probe.

### Tunneling and Puncture (Subcutaneous)

Cutting, tunneling, and puncture were qualitatively evaluated by pre-clinical implanters with significant experience performing animal and/or cadaver implant studies. DA slabs of subcutaneous tissue (muscle and fat combination) were evaluated to assess initial incision/cutting, blunt dissection, tunneling, and puncture. Cutting was evaluated with an incision made by an 11-blade scalpel, commonly used for similar procedures. Tunneling and puncture were evaluated using a rigid metal tunneling tool. Figure 7 shows the various tests being performed. Evaluators were asked to rate the nine subcutaneous tissue structures (Subcutaneous Tissue 1 – Subcutaneous Tissue 9) as red, yellow, or green for the three characteristics.

**Figure 7.**
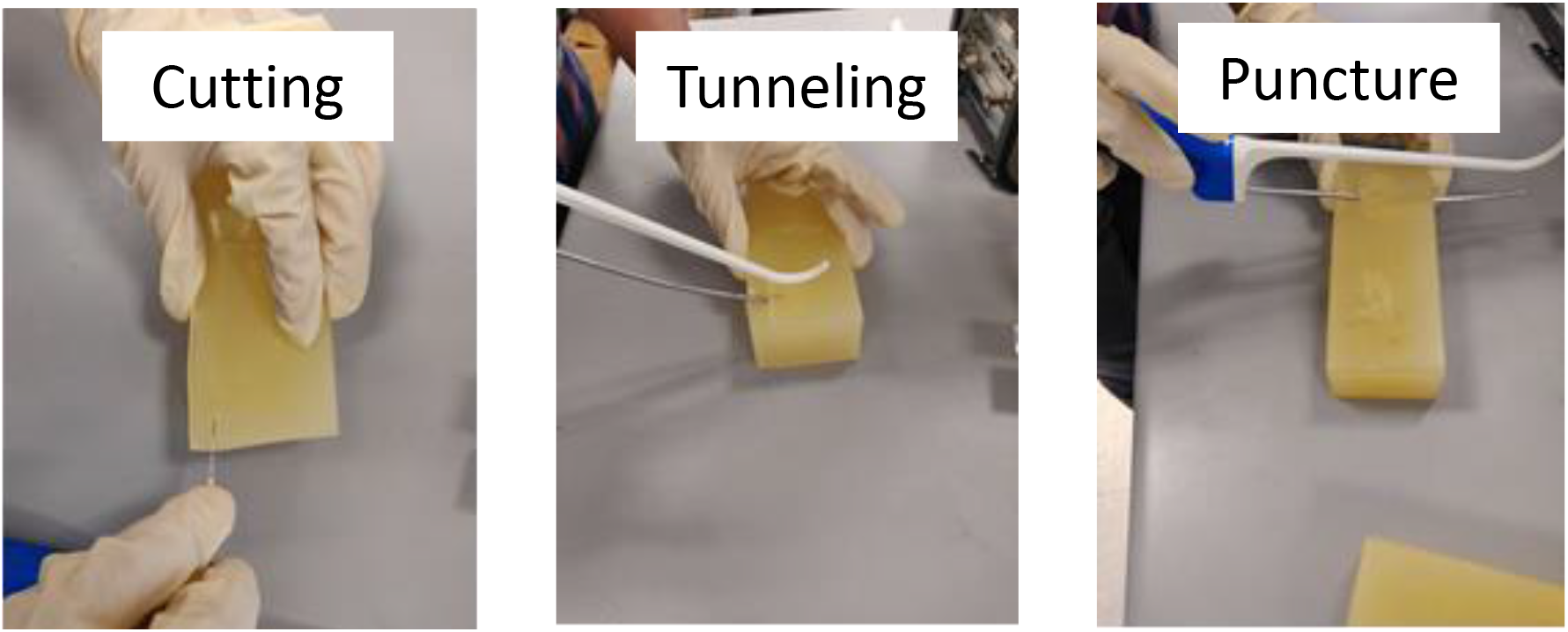
(Left): Incision and cutting with 11-blade scalpel. (Middle): Tunneling with implant tool. (Right): Puncture through the sample wall with implant tool.

## Results

### Stiffness Testing

#### Porcine Liver

Six porcine livers were dissected into 140 samples, a combination from all four lobes, and evaluated for stiffness from 1 to 3 N. Due to the high variability of animal tissue samples, the samples were not separated out based on type of lobe; instead, all of the samples were analyzed together. Figure 8 shows the stiffness values plotted against sample thickness.

**Figure 8.**
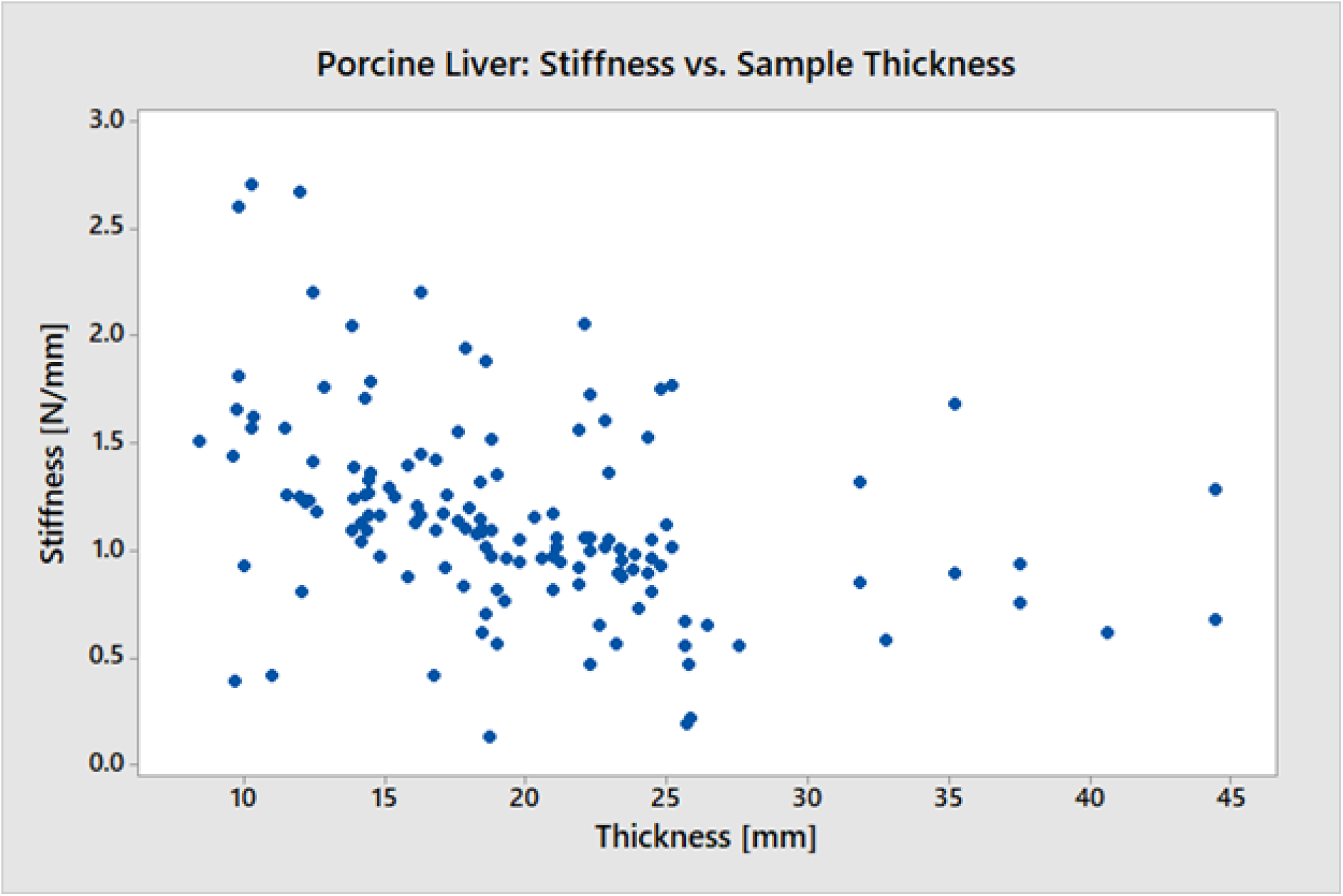
Stiffness (N/mm) plotted against sample thickness (mm) evaluated from 1 to 3 N for all 140 samples harvested from six porcine livers.

#### Digital Anatomy Printed Material

Filter testing was performed on an initial batch of fifteen different experimental DA SO configurations, considered iteration one. The Liver 2 configuration from iteration one was within the range of porcine tissue stiffness values for the two printed thicknesses (15 mm and 25 mm), thus further replicate testing (n = 6) was conducted and a second iteration of different liver configurations was tested. Samples were further tested to identify any differences based on the thickness of the Agilus shell. The stiffness values for iteration one Liver 2 configurations are shown in Figure 9. Both thickness groups (15 mm and 25 mm) demonstrate the same stiffness value trend where the 0.4 mm Agilus shell samples are less stiff than the 1.0 mm Agilus shell samples. The basic statistics for Iteration 1 – Liver 2 are displayed in Table 2, showing the lower standard deviation, thus higher repeatability, of DA SO Liver 2 compared to porcine liver tissue.

**Figure 9.**
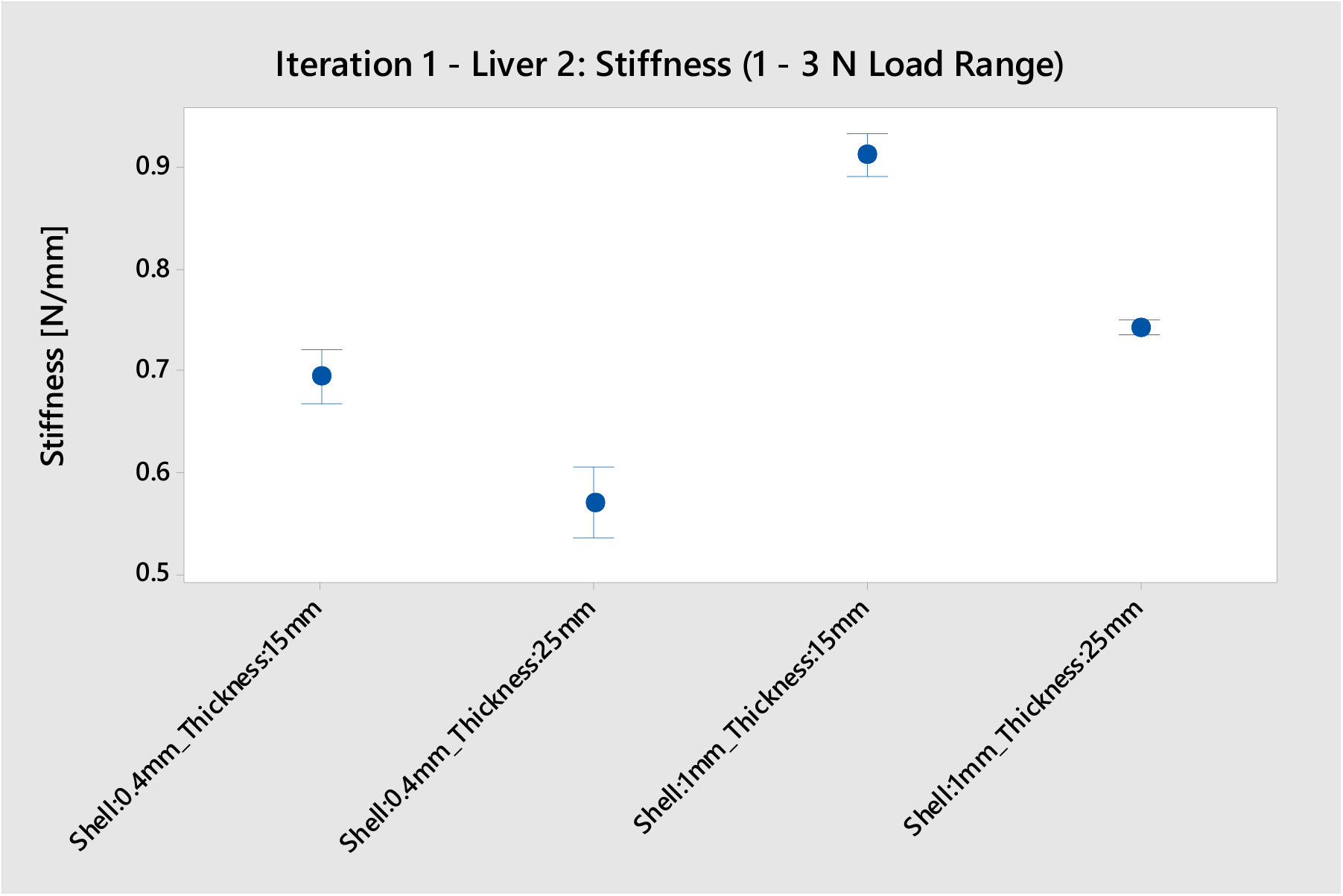
Stiffness values determined from the load vs. extension curve of Iteration 1 - Liver 2 replicates (n = 6) evaluated from 1 - 3 N.

**Table 2.** Basic statistics for the stiffness values determined from the load vs. extension curve of Iteration 1 - Liver 2 replicates (n = 6) and porcine liver tissue (n = 140) evaluated from 1 - 3 N. Printed samples have a lower standard deviation and standard error of the mean, indicating higher reliability than porcine tissue.

| Sample Type | Mean [N/mm] | Standard Deviation | Standard Error of the Mean (SEM) |
| --- | --- | --- | --- |
| Iteration 1 – Liver 2<br>Shell: 0.4 mm \| Thickness: 15 mm | 0.695 | 0.025 | 0.010 |
| Iteration 1 – Liver 2<br>Shell: 0.4 mm \| Thickness: 25 mm | 0.570 | 0.033 | 0.014 |
| Iteration 1 – Liver 2<br>Shell: 1.0 mm \| Thickness: 15 mm | 0.914 | 0.020 | 0.008 |
| Iteration 1 – Liver 2<br>Shell: 1.0 mm \| Thickness: 25 mm | 0.743 | 0.033 | 0.003 |
| Porcine Liver Tissue | 1.144 | 0.455 | 0.038 |

The second iteration of experimental DA SO configurations consisted of seven different variations (Liver 1 – Liver 7). All seven variations were within the range of stiffness values for porcine liver tissue. Figures 10 (Agilus shell: 0.4 mm) and 11 (Agilus shell: 1.0 mm) show the stiffness values of the seven configurations compared to porcine tissue values. All values are within the tissue variation range. For Figure 10, there is a trend across thickness levels where Liver 6 has the highest stiffness and Liver 4 has the lowest stiffness. For Figure 11, the 25 mm thickness samples for Liver 7 and Liver 2 have the lowest and highest stiffness values respectively. Furthermore, the 15 mm thickness samples for Liver 2 and Liver 3 have the lowest and highest stiffness values respectively. The second iteration samples were not tested with replicates, so only general trends were observed.

**Figure 10.**
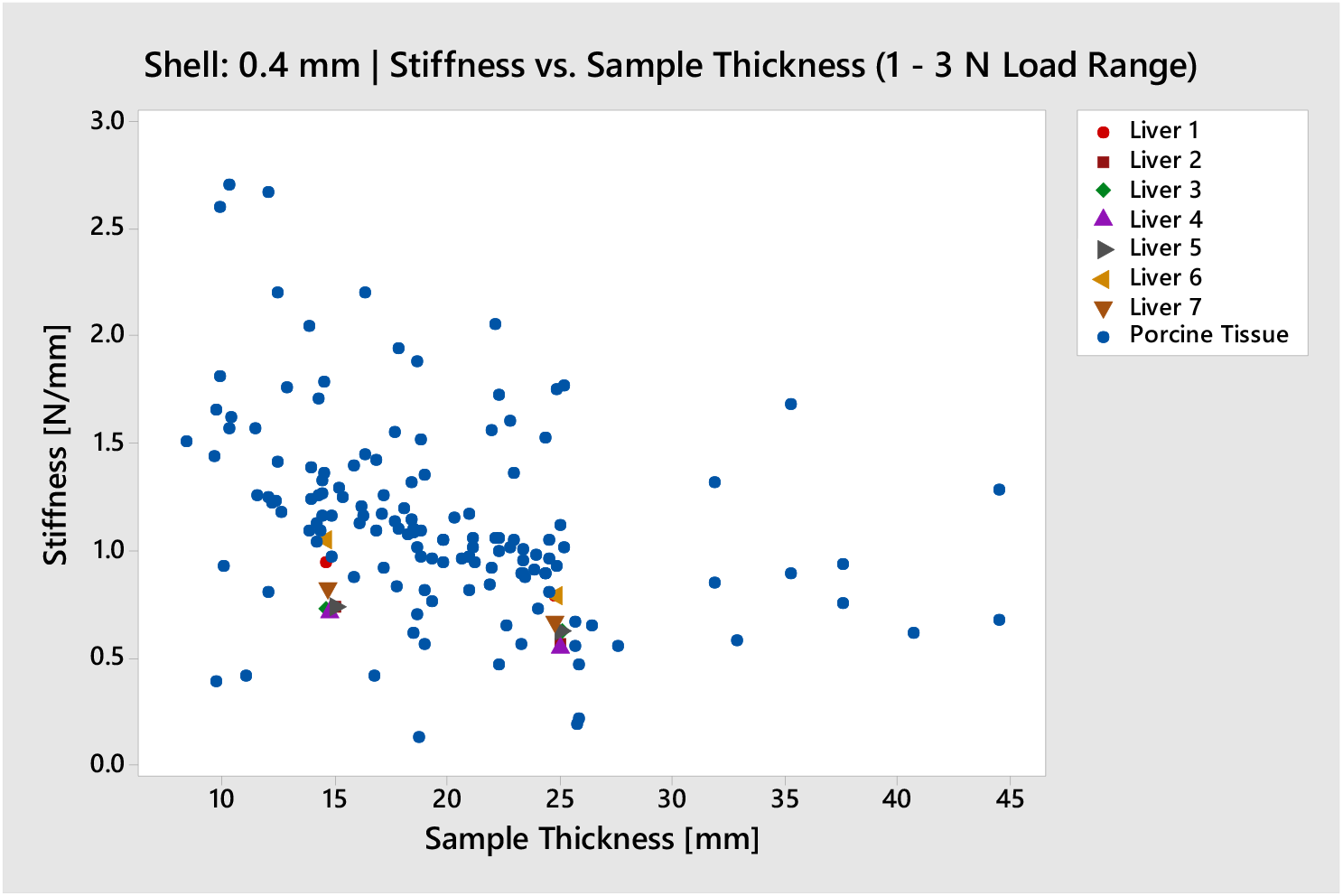
Stiffness vs. Sample Thickness of Iteration 2 DA SO (Liver 1 – Liver 7) with 0.4 mm Agilus shell compared to porcine tissue evaluated from 1 – 3 N. 15 mm and 25 mm: minimum – Liver 4, maximum – Liver 6

**Figure 11.**
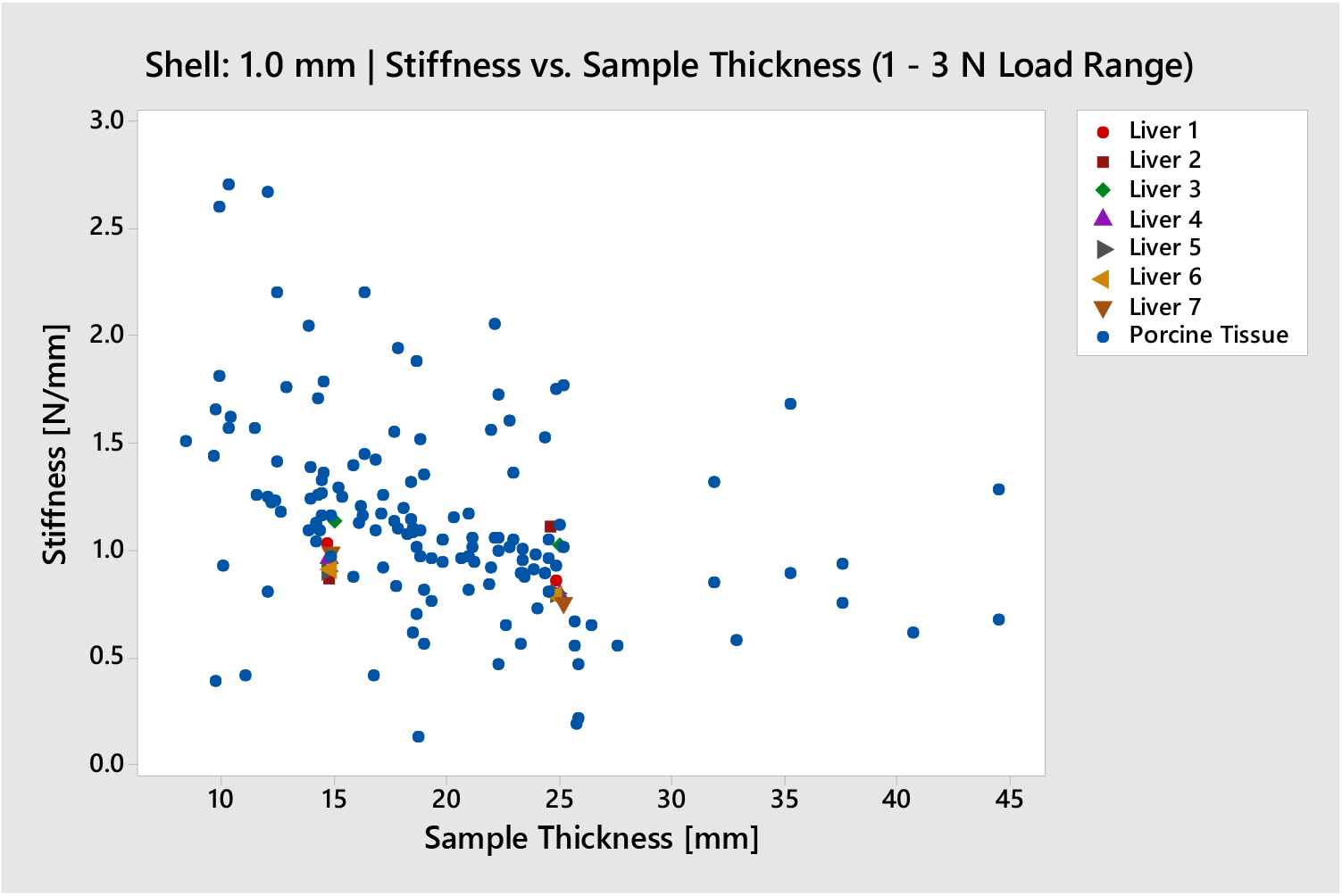
Stiffness vs. Sample Thickness of Iteration 2 DA SO (Liver 1 – Liver 7) with 1.0 mm Agilus shell compared to porcine tissue evaluated from 1 – 3 N. 15 mm: minimum – Liver 2, maximum – Liver 3 | 25 mm: minimum – Liver 7, maximum – Liver 2

### Lubricity Testing

The output analyzed from the lubricity testing was a plot of coefficient of friction vs. time. Due to the sensitivity of the load cell and low load conditions required for non-rigid materials, the raw data was noisy and required processing. The raw data was analyzed from 10 to 50 seconds to allow for steady state, avoiding ramp up and ramp down regions across all test samples. The data was further processed using a simple moving average (SMA) across the entire time interval. Representative plots of the raw and processed data are shown in Figure 12. The measured coefficient of friction values were generated from the boundary conditions used specifically for this lubricity test setup.

**Figure 12.**
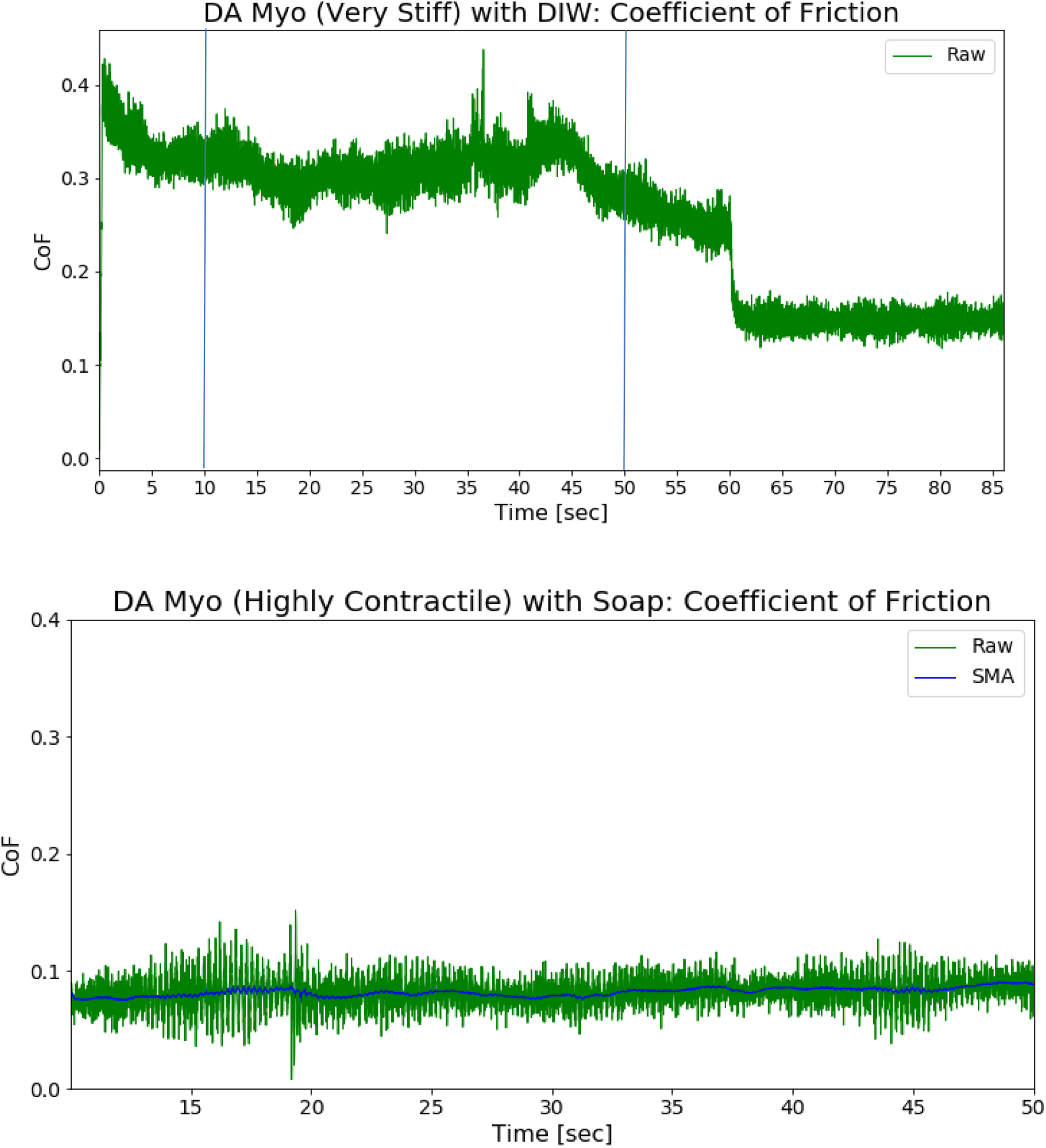
(Top): Raw Coefficient of Friction (Green) vs. Time for DA Myo (Very Stiff) with DI Water over the entire test duration. (Bottom): Processed Coefficient of Friction (Blue) vs. Time plot for DA Myo (Highly Contractile) configuration with a surface treatment of soap.

#### Porcine Epicardium and Aorta

Three porcine hearts were used to generate 33 sample runs on epicardial tissue and 19 sample runs on aortic tissue. Figure 13 shows the grouped runs for both types of tissues. Figure 14 shows the distribution of all the runs of each type of tissue.

**Figure 13.**
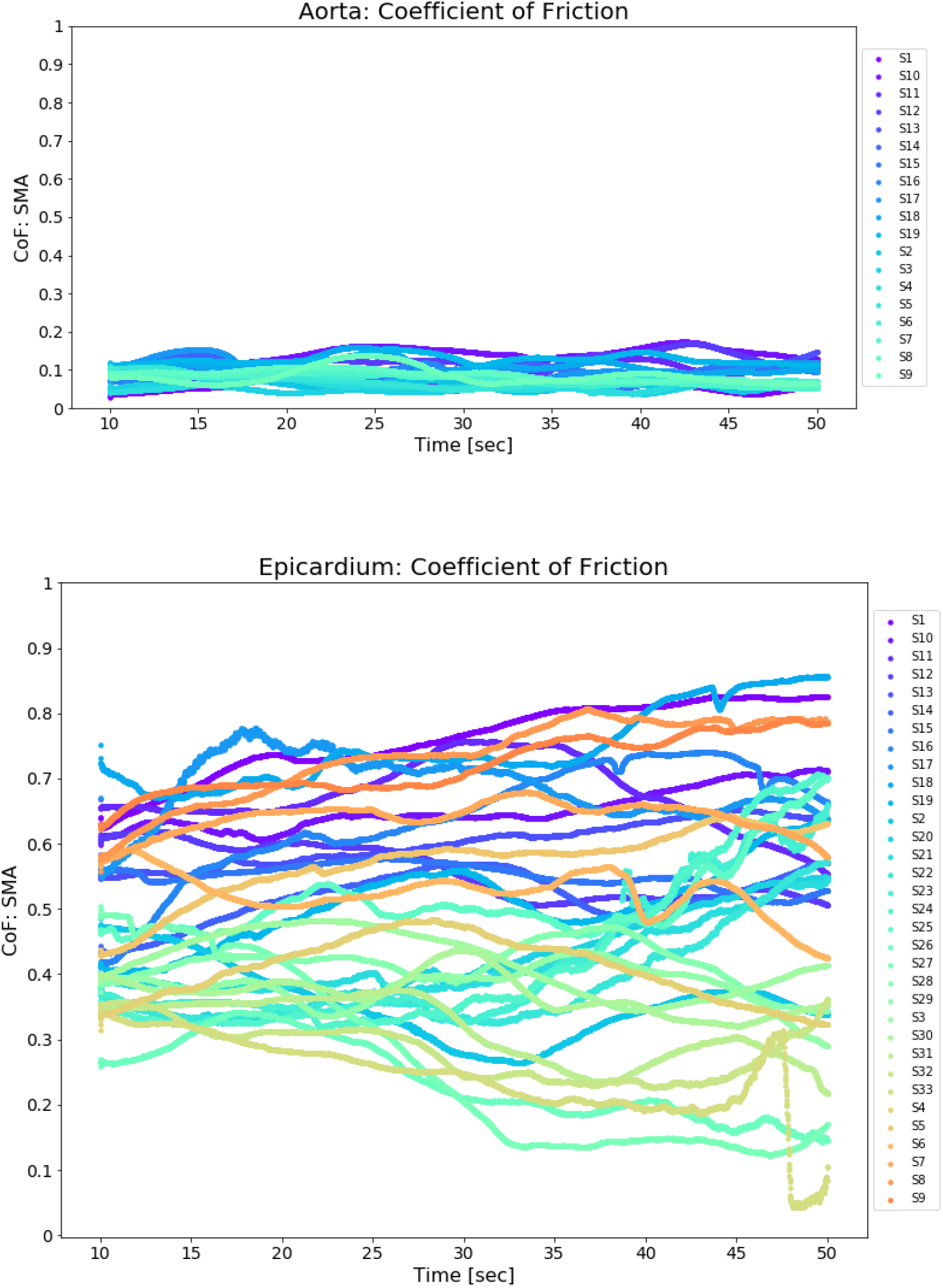
(Top) Coefficient of Friction vs. Time plot for porcine aorta: values are ranging from 0.027 to 0.174. (Bottom) Coefficient of Friction vs. Time plot for porcine epicardium: values are ranging from 0.042 to 0.856.

**Figure 14.**
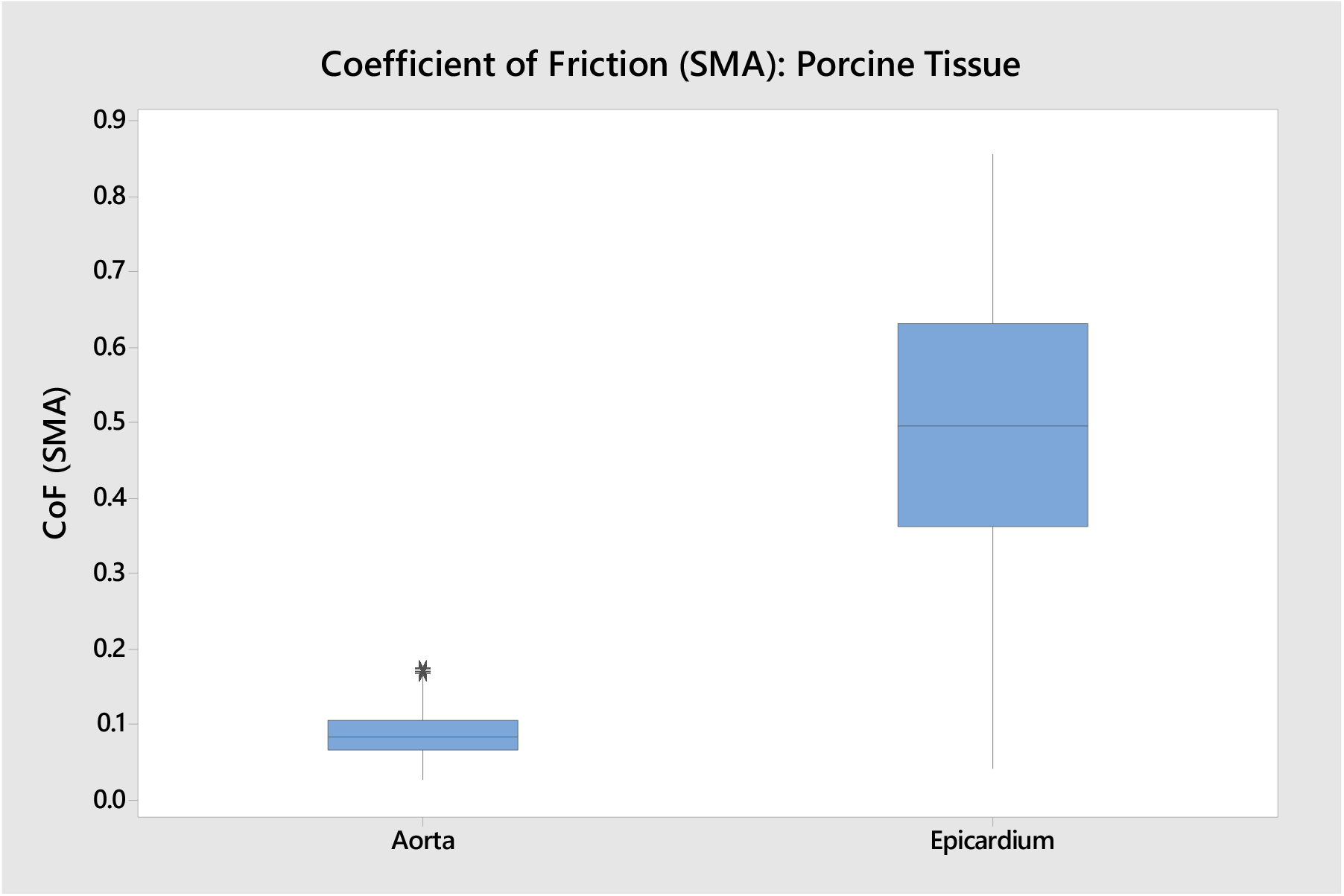
Boxplot of all the runs for epicardium and aorta. Epicardium has a wider spread of coefficient of friction values than aorta.

The basic descriptive statistics for the combined runs of porcine epicardium and aorta are shown in Table 3. The standard deviation of the aortic tissue is much lower than epicardial tissue, and the average coefficient of friction of aortic tissue is 0.088 compared to the average coefficient of friction of epicardial tissue at 0.496. The surface topography from vasculature most likely contributed to the large variation in measured epicardial coefficient of friction values.

**Table 3.** Basic descriptive statistics for tested porcine aorta and epicardium.

| Tissue Type | Mean | Standard Deviation | Minimum | Maximum |
| --- | --- | --- | --- | --- |
| Aorta | 0.088 | 0.028 | 0.027 | 0.174 |
| Epicardium | 0.496 | 0.164 | 0.042 | 0.856 |

#### Digital Anatomy Printed Material

The coefficient of friction of Agilus, and DA Myo (Highly Contractile, Very Stiff, and Extremely Stiff) was observed under no lubricant (dry), DI water, mineral oil, and soap lubricant layer conditions. For the three lubricant layers, n = 5 tests were conducted per DA configuration including Agilus. For the no lubricant condition, n = 3 tests were conducted to demonstrate the upper bound for the coefficient of friction. Figure 15 shows a representative plot of the simple moving average coefficient of friction for n = 5 runs. Due to the variability present between runs, all runs for each DA configuration and lubricant type were grouped together. Figure 16 shows how the various printed configurations compare with the porcine tissue values.

**Figure 15.**
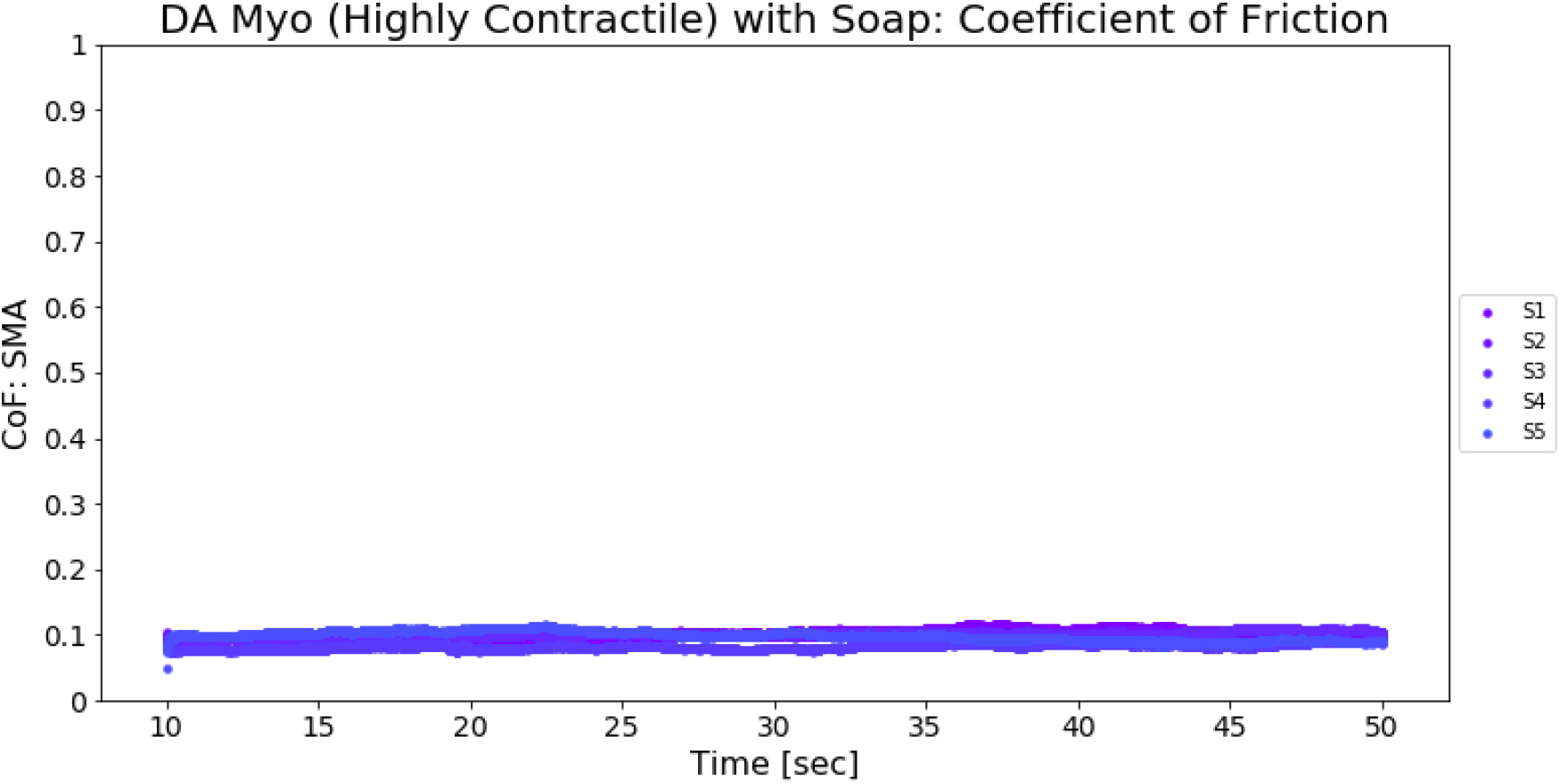
Coefficient of Friction vs. Time plot for 5 samples of DA Myo – Highly Contractile with a surface treatment of soap.

**Figure 16(a-e).**
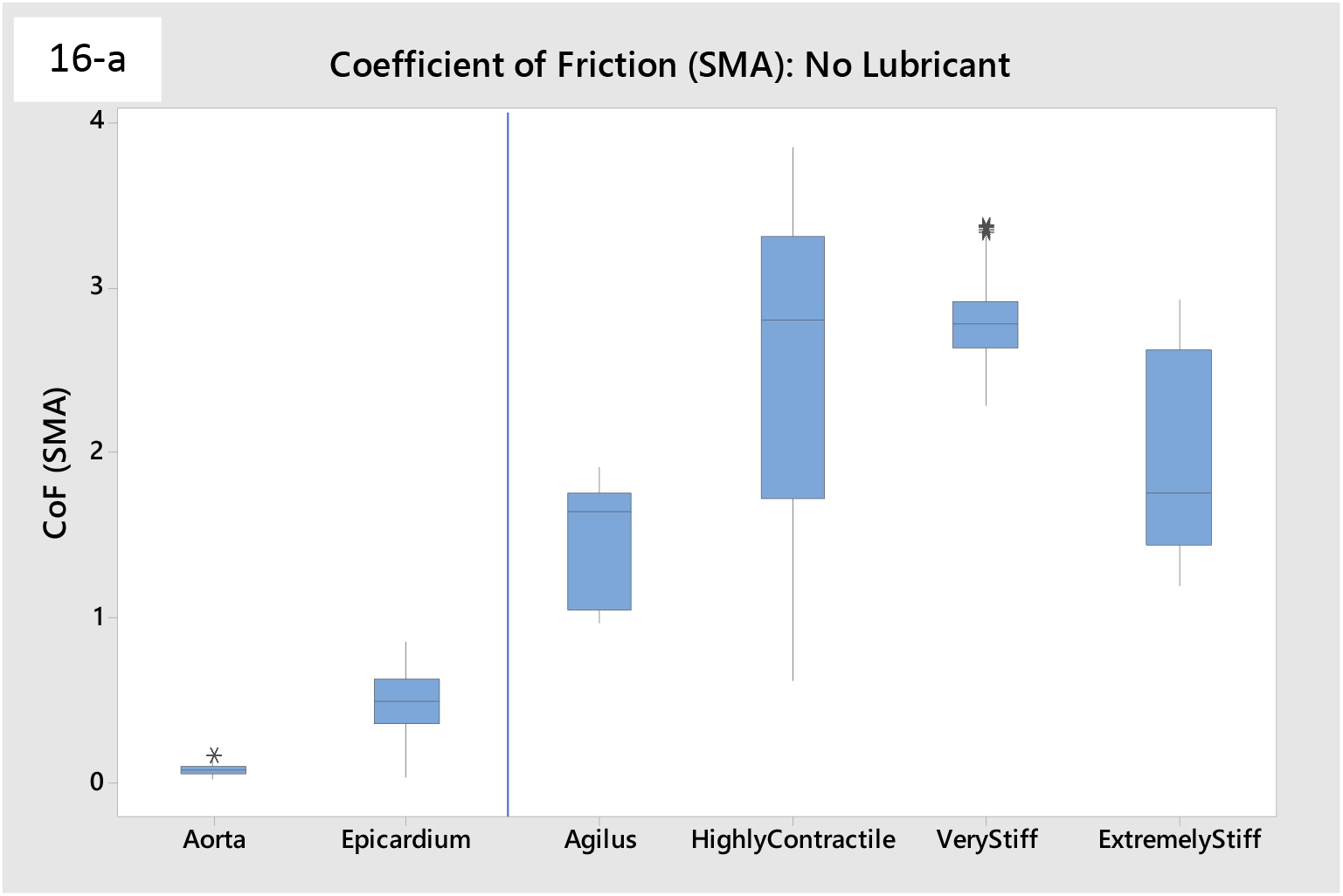
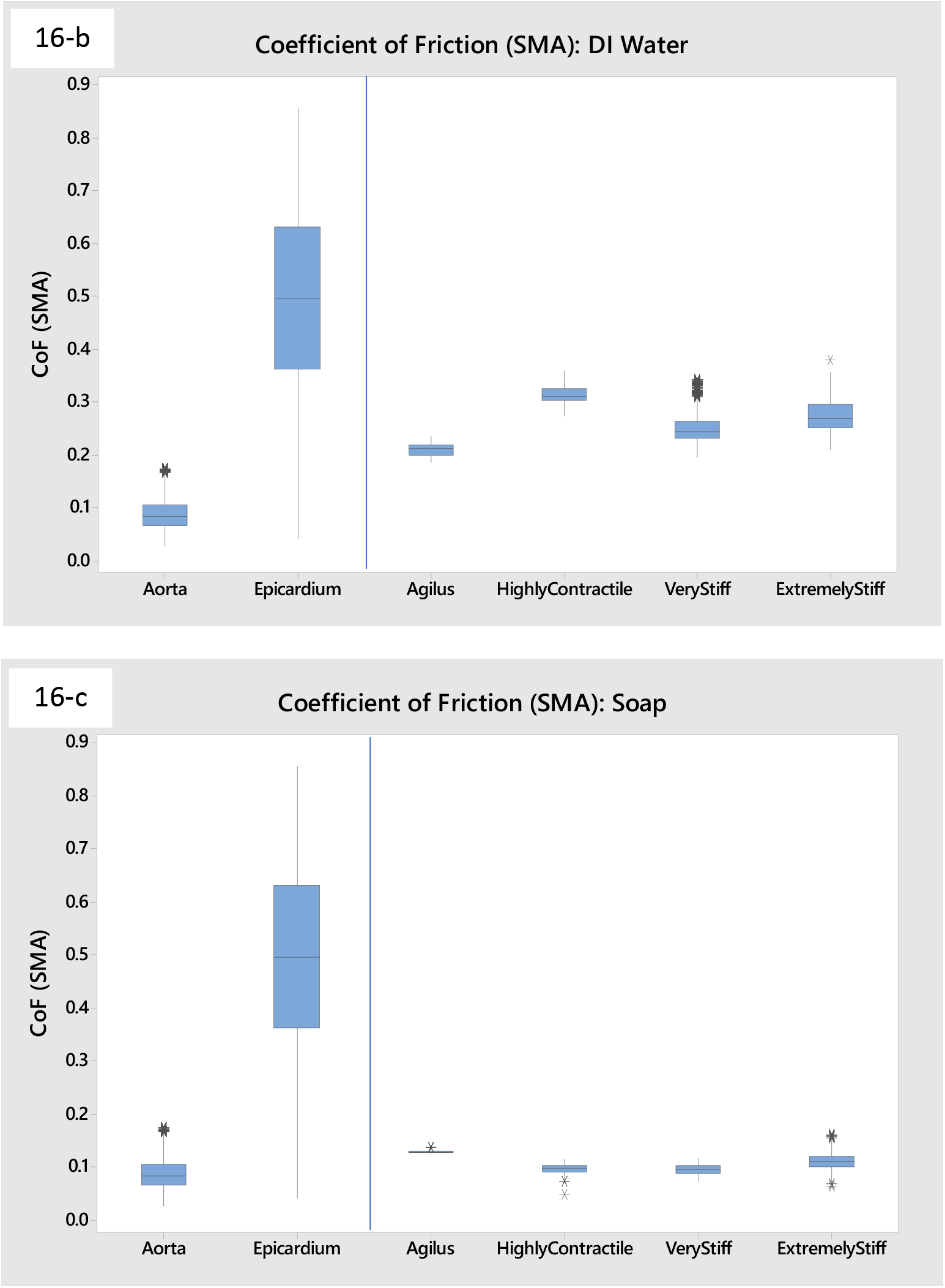
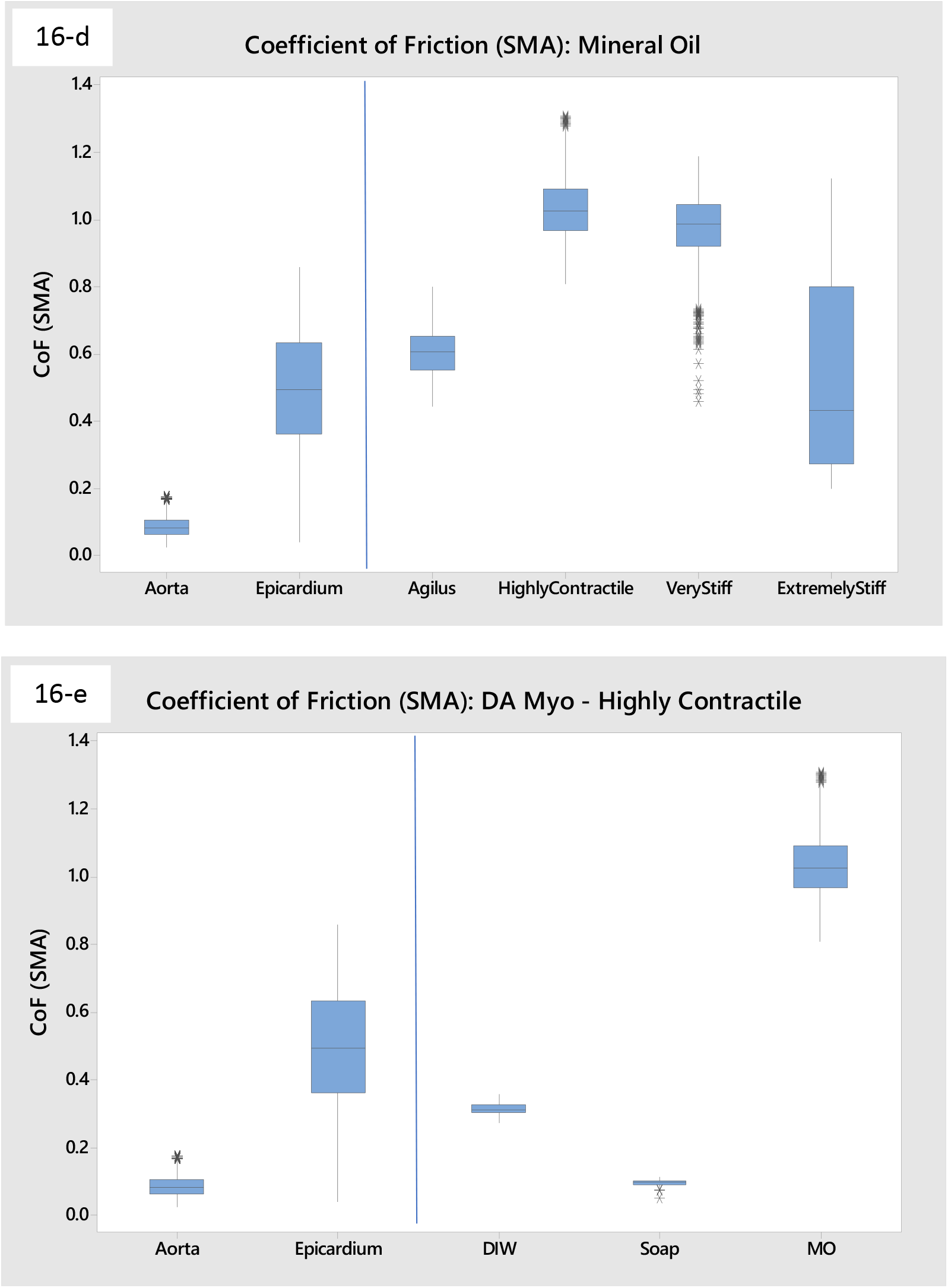
Boxplots of the various DA configurations along with the different lubricant layers are shown in comparison to porcine tissue coefficient of friction values. Fig. 16-e depicts the effect of the three lubricant layers (DI water, soap, and mineral oil) on the coefficient of friction for DA Myo – Highly Contractile.

The dry condition for all of the DA configurations and Agilus resulted in much higher coefficient of friction values than both porcine aorta and epicardium. The printed materials tested with a lubricant layer of DI water had coefficient of friction values between that of porcine aorta and epicardium; the values for the printed materials were within the range found for porcine epicardial tissue. All of the printed materials were close in value to porcine aorta when tested with a lubricant layer of soap, with Agilus being slightly higher in value than the DA materials. There was a distinct difference in coefficient of friction values for printed materials tested with a layer of mineral oil. In general, DA Myo – Highly Contractile and DA Myo – Very Stiff had higher coefficient of friction values than porcine epicardium, but DA Myo – Extremely Stiff had values which encompassed the lower to upper quartile of the porcine epicardium. Agilus also was very close to porcine epicardium under these conditions.

### Qualitative Assessment for Cutting, Tunneling, and Puncture

Cutting, tunneling, and puncture of DA Subcutaneous Tissue material were qualitatively assessed by three pre-clinical implant specialists. Overall, tunneling and puncture for some of the DA Subcutaneous Tissue configurations were close to real tissue behavior while cutting was not. For the favorable configurations, the tunneling of the metal rod through the TissueMatrix provided realistic resistance and the puncture through the Agilus shell provided the tactile feedback that the reviewers expected. In general, the feedback regarding cutting was that too much force was required to create the initial incision and too much drag was present when extending the cutting motion. All three implant specialists were impressed by how some configurations resulted in TissueMatrix realistically mimicking blunt dissection through subcutaneous fat. Overall, all three reviewers had very positive comments and expressed that there were DA Subcutaneous Tissue configurations which were close to functioning as bench level test platforms even with the limitations in cutting. The reviewers indicated that the different configurations were impressive and that there were several configurations that they could use. One reviewer indicated that with further development, DA models such as these could help reduce preliminary animal testing. Figure 17 shows the evaluation of the different DA Subcutaneous Tissue configurations for cutting, subcutaneous fat, and tunneling with puncture.

**Figure 17(a-c).**
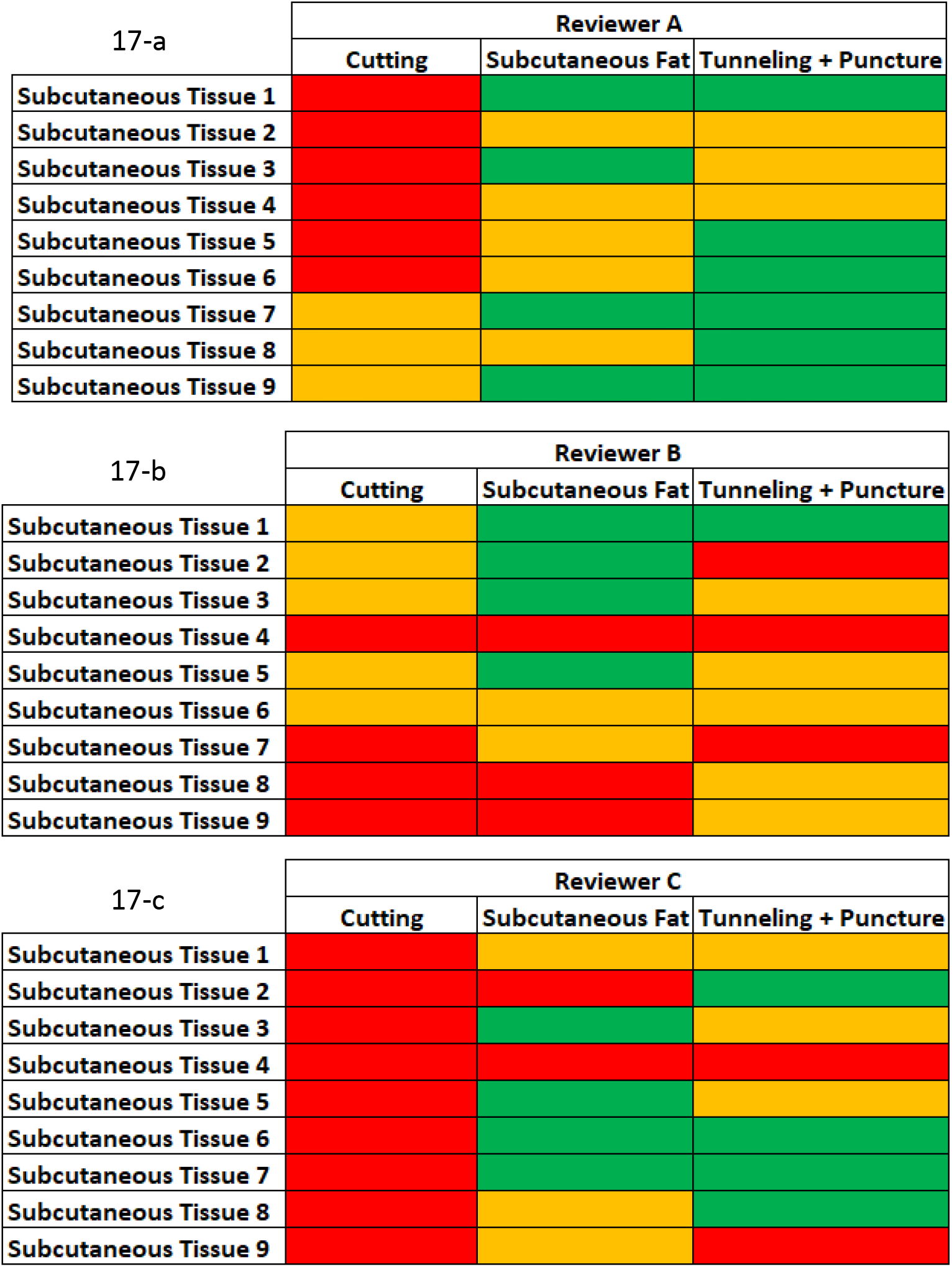
Qualitative evaluation of cutting, subcutaneous fat (tactile response of TissueMatrix), and tunneling (combined with puncture) of Subcutaneous Tissue DA configurations 1 – 9. Red – not realistic; Yellow – Acceptable with Improvements; Green – Acceptable without need for significant improvements

All three reviewers designated multiple DA Subcutaneous Tissue configurations which were acceptable for tactile feedback, tunneling, and puncture. Based on the qualitative feedback, DA Subcutaneous Tissue configurations 1, 3, 5, and 6 show promising results for tactile response of subcutaneous fat, tunneling, and puncture.

## Discussion

This paper has evaluated the stiffness and lubricity of 3D printed DA models and compared to those of porcine tissue. Liver was selected as the soft organ to be simulated using DA. The elastic modulus of porcine liver is non-linear [9,13], the stiffness being higher at larger displacements, whereas the 3D printed DA specimens are more linear. Simulating the low stiffness region using 3D printing is expected to be more challenging than higher stiffness. Hence, the stiffness of the 3D printed samples, and the porcine liver tissue were compared for the low force region of 1 – 3 N, which corresponds to a low displacement and lower stiffness. The ability to 3D print DA models with the stiffness of liver tissue, allows for the potential use of such models for simulating implant procedures and training.

The experimental DA SO liver configurations show promise in their ability to match porcine liver tissue stiffness. While some of the digital materials were evaluated to be on the lower range of tissue stiffness values, the different configurations for the DA SO Liver iterations are within the average stiffness values for tissue at the given thicknesses. Additionally, adjustments in sample thickness could be used to further adjust the stiffness value of a model. Furthermore, the stiffness of 3D printed parts are much more consistent between samples compared to that of porcine tissue, which is highly variable between specimens. Similarly, the 3D printed DA SO liver configurations were consistent with respect to stiffness matching the compliance of biological liver tissue. The repeatability of the material, combined with its tissue realistic stiffness values can provide a significant advantage to 3D printed models as alternatives to preclinical animal or cadaver testing for certain applications; however, there are limitations in the current DA materials, particularly in relation to the non-linear elastic modulus properties. The DA configurations could be adjusted to simulate the higher stiffness at higher displacements, but designing a 3D printed DA configuration to simulate both the low stiffness at low displacements, and high stiffness at high displacement may require advancements in material technology for 3D printing.

Another important aspect of 3D printed models is the lubricity of the models, and replicating in vivo conditions, and haptics requires adjustment of the surface coefficient of friction of the model. For the Agilus and DA Myo configurations, there are commonly available surface treatments which were able to match the coefficient of friction of the printed material to porcine tissue. Furthermore, the lubricant layers formed by the different surface treatments allowed for some flexibility to match the coefficient of friction of porcine epicardium or aortic intima. By combining the ability to modify the coefficient of friction with tissue realistic stiffness values, DA materials can allow for more appropriate bench testing boundary conditions prior to utilizing animal or cadaver models.

The experimental DA Subcutaneous Tissue configurations are promising when evaluated in terms of qualitative tactile feedback. Overall, there was positive feedback in terms of the procedural likeness of DA materials compared qualitatively to animal and cadaver models for puncture and tunneling through the subcutaneous models. There were, however, limitations in the simulation of cutting of these configurations. This feedback matches previous evaluations for cutting, and suture retentions strength [12]. The peak suture pull force to failure, which is similar in mechanism to cutting, was reported to be much lower for 3D printed DA samples compared to porcine myocardium. A similar low suture retention strength was reported for 3D printed alginate structure [14]. Although there are methods to increase the suture holding force, they may result in an increase in stiffness and/or deterioration in puncture or cutting. More work is needed, including improvements in materials, to address these current limitations of 3D printed DA materials.

The versatility to mimic the mechanical properties of biological tissue, as well as the qualitative haptics can allow DA materials to be appropriately used to create anatomical models for both quantitative benchtop testing, as well as for implant procedure planning and training purposes. The methodology and testing used in this work can also be used for further development and evaluation of new material configurations.

## Notes

### Competing Interest Statement

The experimental 3D printed samples tested in this study were printed and provided by Stratasys, Ltd. Additionally, a modest discount on the J750 - Digital Anatomy printer and printer materials for testing were provided by Stratasys, Ltd.

